# Identification of stable reference genes for quantitative real-time PCR in human fibroblasts from lymph nodes and synovium

**DOI:** 10.1101/2024.07.12.603332

**Authors:** S. Rasouli, C.M.J. van Ansenwoude, J.F. Semmelink, L.G.M. van Baarsen, T.A. de Jong

**Author notes:** Corresponding author: Lisa G.M. van Baarsen. Shared final author (L.G.M. van Baarsen, T.A. de Jong).

## Abstract

Real-time quantitative PCR (RT–qPCR) has emerged as an accurate and widely used technique for measuring gene expression levels. However, its reliability depends on the selection of appropriate reference genes to normalize for sample input. Accordingly, the identification of reference genes characterized by stable expression in cells and conditions of interest is essential for ensuring accurate expression values. To date, no study has specifically identified suitable reference genes for primary human cultured fibroblast-like synoviocytes (FLS) and lymph node stromal cells (LNSCs) within the context of rheumatoid arthritis (RA). These stromal cells play a critical role in the pathogenesis of disease. In this study, we evaluated the suitability of 15 candidate reference genes for normalizing transcript expression in FLS and LNSCs subjected to various in vitro stimuli. We included traditional reference genes often used for transcript normalization in fibroblasts as well as candidate genes identified as suitable reference genes via GeneVestigator analysis of publicly available transcriptomic data. RefFinder algorithms were used to identify the most stable reference genes for transcript normalization across the cell types and different experimental conditions. We determined that the optimal number of reference genes for every experimental condition tested was two; *RPLP0* and *POLR2G* exhibited the greatest stability across different experimental conditions for LNSCs. However, for FLS, we observed greater variability in the most stable reference genes across different experimental conditions. Although *POLR2G* and *TBP* emerged as the most stable reference genes under unstimulated conditions, our findings indicated that FLS require distinct reference genes for transcript normalization depending on the specific experimental conditions. Validation of the selected reference genes for normalizing the expression levels of metabolic genes in unstimulated FLS emphasized the importance of prior evaluation of potential reference genes, as arbitrary selection of reference genes could lead to data misinterpretation. This study constitutes the first systematic analysis for selecting optimal reference genes for transcript normalization in different types of human fibroblasts. Our findings emphasize the importance of proper selection of reference genes for each experimental condition separately when applying standard quantitative PCR technology for assessing gene expression levels.

## Introduction

Real-time quantitative polymerase chain reaction (RT-qPCR) is one of the most widely used, accurate and sensitive methods for the detection and quantification of mRNA transcripts. The nowadays commonly applied RT-qPCR technology has played a significant role in revolutionizing biology due to its high sensitivity, specificity, high-throughput detection, and versatility (1, 2). Quantitative PCR has overcome the limitations of end-point PCR by studying amplification in real time during the exponential phase, while standard PCR measures amplification during the plateau phase (3). The ability to accurately quantify mRNA transcript levels is invaluable for understanding gene expression patterns and regulatory mechanisms in numerous cellular contexts, providing critical insights into cellular functions and disease mechanisms (4).

In RT-qPCR, the selection of reference genes, also known as “housekeeping genes,” is critically important for accurate gene expression analysis. These genes are constitutively expressed across various tissues and under various experimental conditions, making them ideal internal controls for transcript normalization (5). These internal controls are used to account for sample-to-sample biases, which can result from variations in total RNA content, RNA stability, enzymatic efficiencies or discrepancies in sample loading. Proper normalization using reliable reference genes corrects for these technical variations and ensures that the measured gene expression levels are accurate, consistent, and comparable across different samples (6). Without the use of validated reference genes, the results of RT-qPCR could be misleading, emphasizing the pivotal role they play in the accuracy and reliability of gene expression studies.

Furthermore, to serve as a reliable reference, the expression of reference genes should remain unaltered regardless of the experimental conditions (3). Two primary challenges affecting the use of reference genes in RT-qPCR quantification and normalization are fluctuations in reference gene stability across different experimental settings and technical errors due to the relatively strong expression of some reference genes, which often differ by several orders of magnitude from the target gene transcripts. In the normalization process, these factors may introduce significant bias, leading to potential misinterpretation of data, especially when relying on a single gene for normalization, a preferred approach for many researchers (5).

Many researchers select commonly used reference genes without verifying their stability across different experimental conditions. For decades, the field of autoimmunity, including rheumatoid arthritis (RA), has focused primarily on cells from the hematopoietic immune system. However, recent studies have revealed the significant role of fibroblasts in disease pathogenesis (7), highlighting the necessity of reassessing the suitability of reference genes when investigating diverse cell types. Despite the frequent use of assumed stable reference genes such as *GAPDH* (cellular metabolic pathway member) and *ACTB* (cytoskeleton component), research has shown significant variations in the expression levels of these genes under different experimental conditions, emphasizing the need for reliable reference genes (5, 8). The current research addresses a crucial gap in understanding gene expression dynamics in RA-associated fibroblasts, enhancing the accuracy and reliability of RT-qPCR normalization in future studies.

RA is an inflammatory autoimmune disease primarily affecting synovial joints and is characterized by chronic synovial inflammation (9-12). Understanding the early phases of RA, before clinical arthritis symptoms appear, could aid in developing preventive strategies. Insights into the disease’s earliest stages revealed the presence of circulating autoantibodies such as rheumatoid factor (RF) and anti-citrullinated peptide antibodies (ACPA) in peripheral blood years before clinical manifestation (RA-risk phase) (11, 13), indicating systemic autoimmunity preceding synovial inflammation. The pathogenesis of RA involves a complex interplay between immune dysregulation and stromal microenvironment alterations, prominently involving fibroblast-like synoviocytes (FLS) (14-16) and lymph node stromal cells (LNSCs) (17, 18). In RA, high levels of inflammatory mediators are produced that perpetuate inflammation, which could lead to joint damage and tissue destruction (19-21). Additionally, LNSCs, which are integral to lymph node architecture and function, play a pivotal role in immune homeostasis and tolerance induction (22-26). Efforts to elucidate the molecular mechanisms underlying FLS and LNSC dysfunction hold promise for advancing precision medicine approaches in RA (27).

To date, no studies have systematically validated candidate reference genes for transcript normalization in human FLS and LNSC populations isolated from patients and controls. Given the pivotal role of FLS in RA pathogenesis, identifying reliable reference genes in fibroblasts is crucial for accurate gene expression analysis. Our objective was to identify and validate the most suitable and stable candidate reference genes for accurate data normalization when performing RT-qPCR in human LNSCs and FLS under various experimental conditions. This approach is essential for ensuring robust and accurate quantification of gene expression changes, thereby enhancing our understanding of the molecular mechanisms underlying FLS and LNSC dysfunction in RA pathogenesis.

## Methods

### Study subjects

Individuals with arthralgia and/or a family history of RA who were positive for anti-citrullinated protein antibodies (ACPAs; detected by the anti–cyclic citrullinated peptide [anti–CCP] antibody test (CCPlus anti-cyclic citrullinated peptide 2 ELISA (ULN 25 kAU/L) Eurodiagnostica, Nijmegen, the Netherlands) and without any evidence of arthritis upon examination were included. These individuals were considered to be at risk of developing RA (RA-risk individuals), characterized by the presence of systemic autoimmunity associated with RA, but without clinical arthritis (defined as phase c+d, according to EULAR recommendations) (9, 28). Additionally, we included patients with rheumatoid arthritis (ACR/EULAR 2010 criteria (28), osteoarthritis patients (OA) and seronegative controls without any history of autoimmunity and inflammatory disease and no present or previous use of disease-modifying anti-rheumatic drugs (DMARDs) or biologicals. Study subjects were recruited either via the outpatient clinic of the Department of Rheumatology and Clinical Immunology at the Amsterdam UMC, via referral from the rheumatology outpatient clinic of Reade, Amsterdam, or by testing family members of RA patients in the outpatient clinic. Lymph node (LN) tissues were collected by ultrasound-guided inguinal LN core needle biopsy as previously described (29). Synovial tissues were collected during mini-arthroscopic synovial tissue sampling of a knee joint at baseline (RA-risk and RA patients (30, 31)) during joint replacement surgery (RA and OA patients) or during orthopedic surgery of the knee for e.g., ligament rupture (seronegative controls). The study was performed according to the principles of the Declaration of Helsinki (32), approved by the Institutional Review Board of the Amsterdam UMC and all study subjects gave their written informed consent.

### Cell culture and experimental conditions

#### Stromal cell culture, senescence induction and dasatinib treatment

LNSCs and FLS were isolated and expanded *in vitro* as previously described, generating cultures containing lymph node fibroblasts (25) or synovial fibroblasts, respectively (33). Complete cell culture medium consisted of Dulbecco’s Modified Eagle Medium (DMEM), low glucose (Gibco, Bleiswijk, The Netherlands) supplemented with 1% penicillin/streptomycin (10,000 U/mL, Gibco), 10 mM 4-(2-hydroxyethyl)-1-piperazineethanesulfonic acid (HEPES) buffer (Gibco) and 10% fetal bovine serum (FBS) (Biowest, Nuaillé, France). Cells were harvested using 0.05% trypsin/5 mM ethylenediaminetetraacetic acid (EDTA; Thermo Fisher Scientific, Landsmeer, The Netherlands) and counted using trypan blue (Sigma–Aldrich, Zwijndrecht, The Netherlands) in a Burker Turk hemocytometer (VWR, Amsterdam, The Netherlands).

To induce senescence, LNSCs were irradiated in culture flasks at 10 Gy using CellRad+ (Precision X-ray, Madison, CT) and passaged simultaneously with nonirradiated cells from the same donor. To remove senescent cells, LNSCs were treated with 5 µM dasatinib (MedChemExpress, Monmouth Junction, NJ) for 24 hours. FLS were irradiated at 5 Gy and treated with 0.25 µM dasatinib for 24 hours, as these cells were more sensitive to irradiation and senolytic treatment. After 24 hours of treatment, the culture flasks were washed with complete cell culture medium and cultured until they reached 80% confluence, after which all the cells were simultaneously harvested and collected in RNA lysis buffer (RLT + beta-mercaptoethanol) (Qiagen, Venlo, The Netherlands).

#### TNFα, LTαβ and IFNγ stimulation

Human fibroblasts were seeded in a 24-well plate and stimulated with tumor necrosis factor-α (TNF-α) (5 ng/ml; Life Technologies, Landsmeer, the Netherlands) plus lymphotoxin α1β2 (50 ng/ml; R&D Systems, Abingdon, UK) or with IFNγ (50 ng/mL, 34-8319, eBioscience, Vienna, Austria) for 24 or 72 hours. After these timepoints, the cells were harvested and collected in RNA lysis buffer.

#### Adipogenesis differentiation

Human fibroblasts were seeded in 12-well plates in adipogenesis differentiation medium consisting of ɑMEM (Lonza, Geleen, The Netherlands) supplemented with 10% FBS, 5 µg/mL insulin, 500 µM IBMX, 1 μM dexamethasone and 1 μM indomethacin (all from Sigma–Aldrich) for 14 days. After 14 days of culture in adipogenesis differentiation medium, the cells were harvested and collected in RNA lysis buffer.

#### Primer design

Sequence information of the candidate reference genes was obtained from the human transcriptome in the NCBI database (https://www.ncbi.nlm.nih.gov/gene/), and gene-specific primers used for RT–qPCR were designed using Primer Blast (Primer designing tool (nih.gov)), PCR Primer Stats (http://www.bioinformatics.org/sms2/pcr_primer_stats.html) and the OligoAnalyzer™ tool (IDT, https://eu.idtdna.com/calc/analyzer). Primers were designed using the Primer-BLAST design tool from the National Library of Medicine with the following parameters: Tm approximately 60°C, amplicon length of 90 to 120 base pairs, yielding primer sequences with an optimal length of 20 nucleotides, and a GC content of 45 to 60%. Whenever possible, primers were designed to allow the amplification of transcript isoforms from all human genotypes. The specificity of the resulting primer pair sequences was validated against the *Homo sapiens* transcript database, and amplicon specificity was confirmed through melting curve analysis. To analyze the amplification efficiency of each primer pair, 10-fold serial dilutions of arbitrary calibrator samples were generated and utilized as templates for RT-qPCR analysis. The standard regression lines, coefficient of determination (R^2^) and amplification efficiency of each primer pair were evaluated using QuantStudio™ Design&Analysis software v1.4.3 (Applied Biosystems, Life Technologies, Zwijndrecht, The Netherlands).

#### Quantitative real-time PCR

Total RNA was isolated using the RNA Micro or AllPrep Kit (Qiagen) according to the manufacturer’s instructions. Subsequently, cDNA was prepared using the RevertAid H Minus First Strand cDNA Synthesis Kit (Thermo Fisher Scientific, Landsmeer, The Netherlands). Quantitative PCR was performed using fast SYBR® Green PCR master mix (Applied Biosystems, Life Technologies, Zwijndrecht, The Netherlands) combined with in-house designed primers (Thermo Fisher). The primer sequences used are listed in Table 1. For detection, we used QuantStudio 3 (Applied Biosystems). An arbitrary calibrator sample was used to correct for interplate differences. To calculate the relative quantity (RQ) of the target genes of interest, the standard curve method was applied.

**Table 1.**
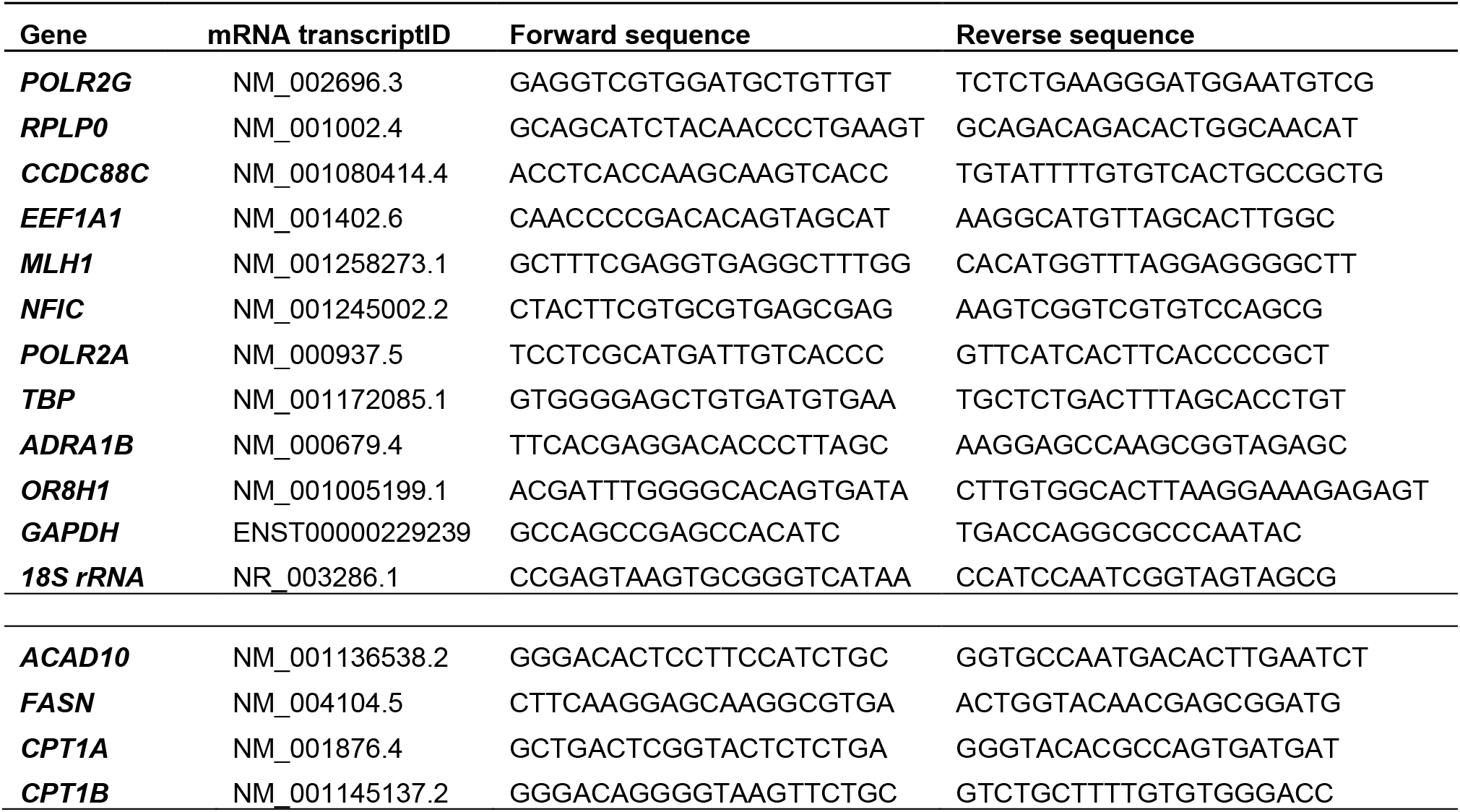
Overview of primer sequences.

RT-qPCR was optimized for each primer pair, and independent biological samples were evaluated for each experimental condition in technical replicates. Melting curve analysis confirmed the presence of a single PCR product from all samples with no primer dimers. The amplification efficiency, ranging from 92 to 98%, was estimated using QuantStudio analysis software. Cycle quantification for each reaction, determined by the maximum point of the second derivative curve, was also calculated using QuantStudio.

#### Evaluation of reference gene expression stability

To identify the optimal reference genes in LNSCs and FLS under different experimental conditions, the expression stability of 12 candidate reference genes was evaluated based on the Ct values obtained from RT-qPCR detection using the delta Ct method, geNorm (integrated in qBase+) and the web-based analysis tool RefFinder (https://blooge.cn/RefFinder/), which analyses and ranks the candidate reference genes using the geNorm, NormFinder and BestKeeper algorithms (34). Among them, geNorm calculates M values, with lower M values indicating greater stability of the reference genes. Moreover, geNorm also analyzes the pairwise variation (V_n_/V_n+1_) between two sequential genes with a suggested cut-off of V < 0.15 for the suitable number of reference genes needed for optimal normalization (34). Based on the weighted ranking of these four methods, RefFinder provides a comprehensive order of the most stable reference genes to the least stable reference genes.

#### Statistical analyses

Statistical analyses were performed using GraphPad Prism version 9.1.0 (GraphPad Software Inc., USA). The data are presented as the mean ± standard error of at least two independent biological replicates for reference gene analyses. Differences in target genes of interest between study groups or experimental conditions are presented as the median ± IQR and were analysed using the Kruskal–Wallis test followed by Dunn’s post hoc test. P-values < 0.05 were considered statistically significant.

## Results

FLS and LNSCs from RA patients, seronegative controls and individuals at risk of developing RA were cultured under different experimental conditions. Fibroblasts were cultured in adipogenesis differentiation medium, stimulated with TNFα LTαβ or IFNγ, or treated with gamma-radiation or senolytics (Figure 1). Potential reference genes were selected based on their common use as reference genes for transcript normalization in fibroblasts, based on their stable expression in transscriptome microarray data, (35-40) or based on relevant literature (Table 2), as described in more detail in the next section (Figure 2).

**Figure 1.**
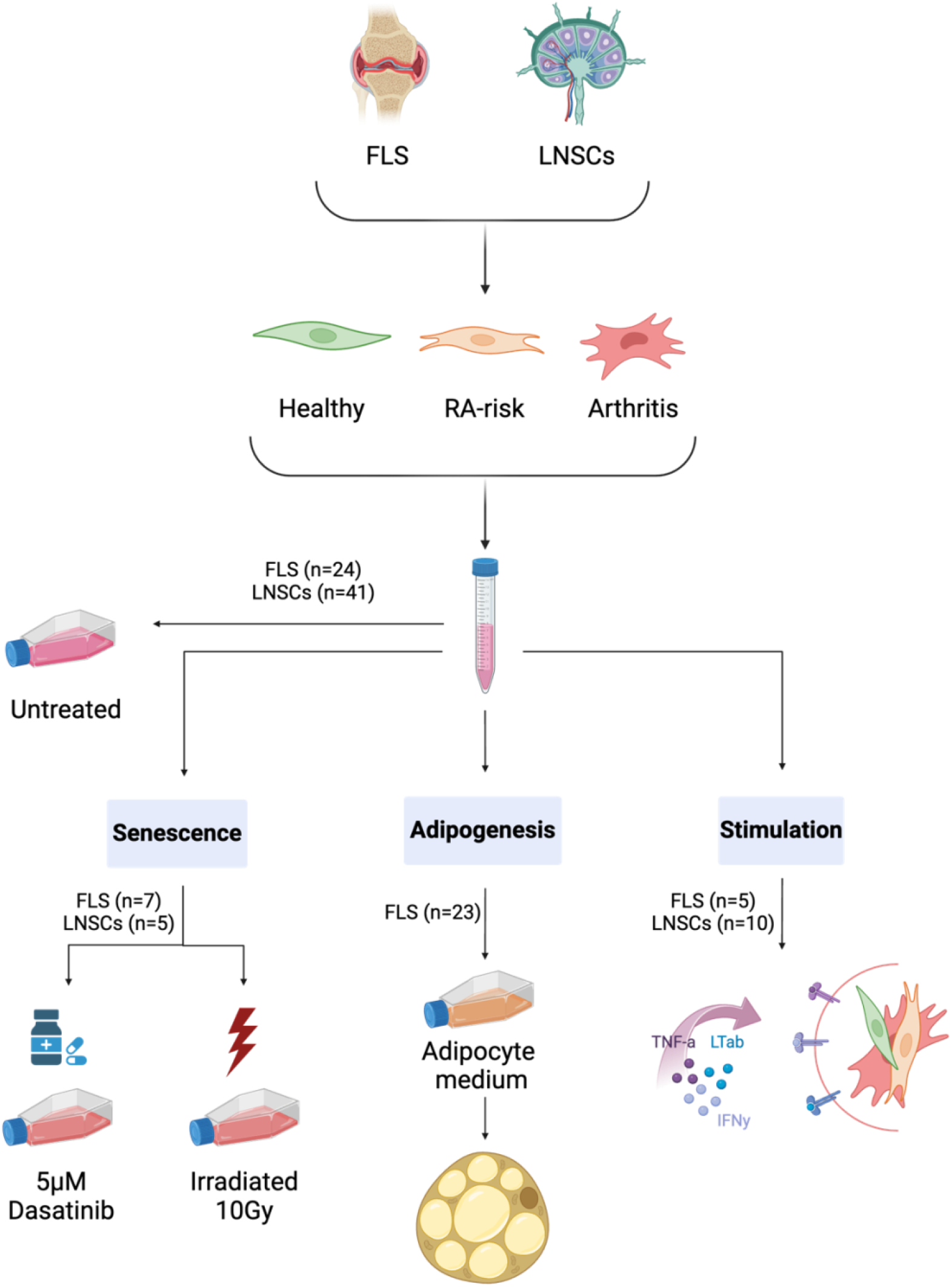
Overview of the experimental conditions. In this study, we used cultured FLS and LNSCs isolated from synovial and lymph node biopsies obtained from healthy individuals, individuals at risk of developing RA, and patients with arthritis. Cells from 41 LNSC and 24 FLS donors were maintained in complete culture medium until they reached 80% confluence and subsequently collected for RNA analyses (untreated). For the senescence experiments, cells from 5 LNSCs and 7 FLS donors were either irradiated or cultured in the presence of dasatinib. Adipogenesis was induced in 23 FLS donors by culturing them in adipocyte medium for 14 days. For the stimulation experiments, cells from 10 LNSCs and 5 FLS donors were treated with IFNy or TNFα in combination with LTαβ.

**Figure 2.**
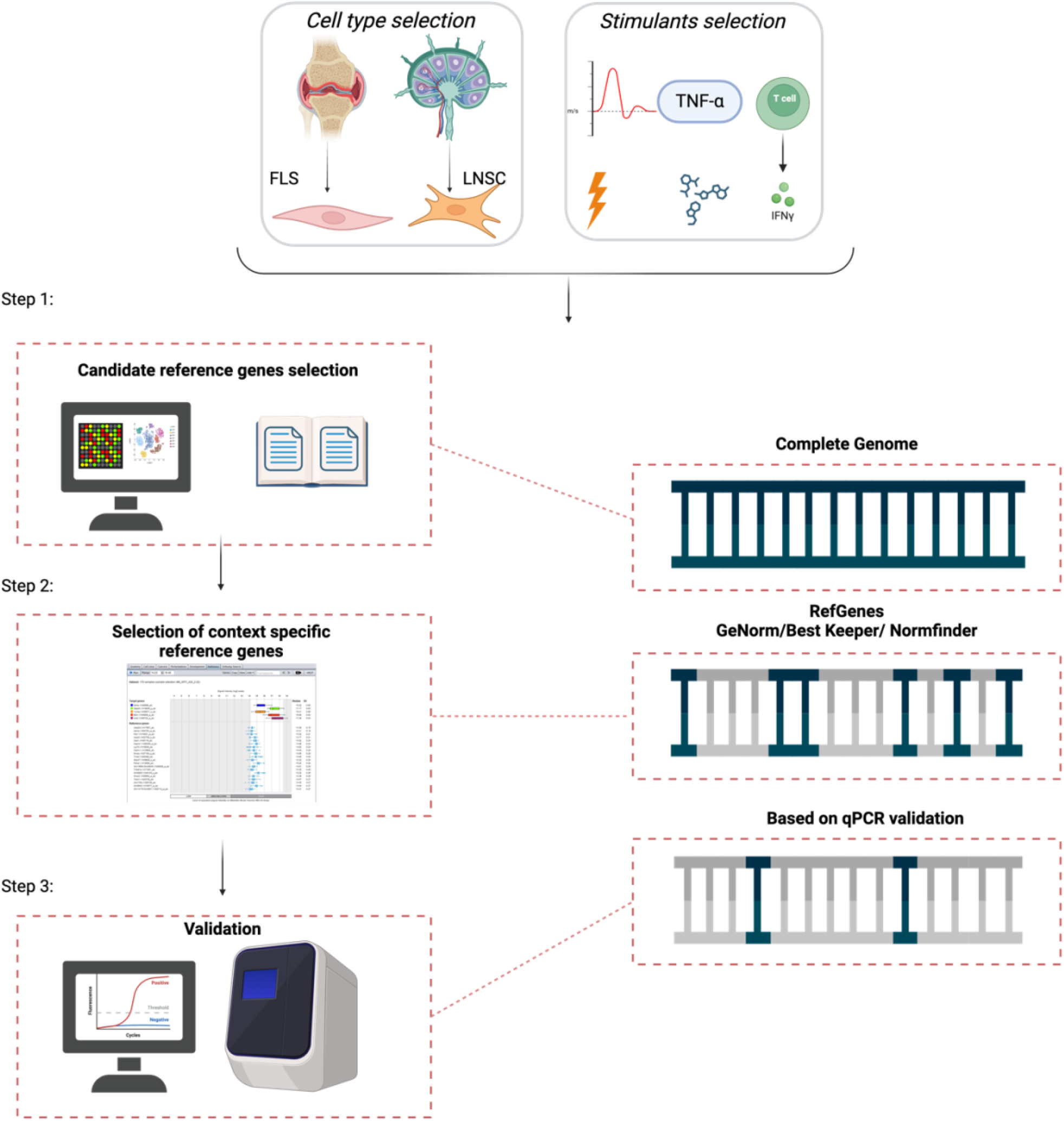
Schematic workflow for the identification of suitable reference genes in FLS and LNSCs. FLS and LNSCs were chosen due to their central role in the pathogenesis of RA. The selected stimulants included adipogenesis, tumor necrosis factor-alpha + lymphotoxin αβ (TNFαLTαβ), interferon-gamma (IFN-γ), irradiation, and dasatinib treatment. **Step 1:** Initial selection of reference gene candidates was conducted via comprehensive genome analysis of publicly available microarray and RNA-seq data and from the literature. **Step 2:** Candidate reference genes were selected based on the context of our study: the genes should not play a role in pathways of interest/related to the study objectives. Candidate reference genes were selected utilizing RefGenes software. **Step 3:** The optimal candidates were identified using GeNorm, NormFinder and BestKeeper tools based on their stable expression levels under our experimental conditions. Finally, the selected reference genes were validated through RT-qPCR analysis.

**Table 2.**
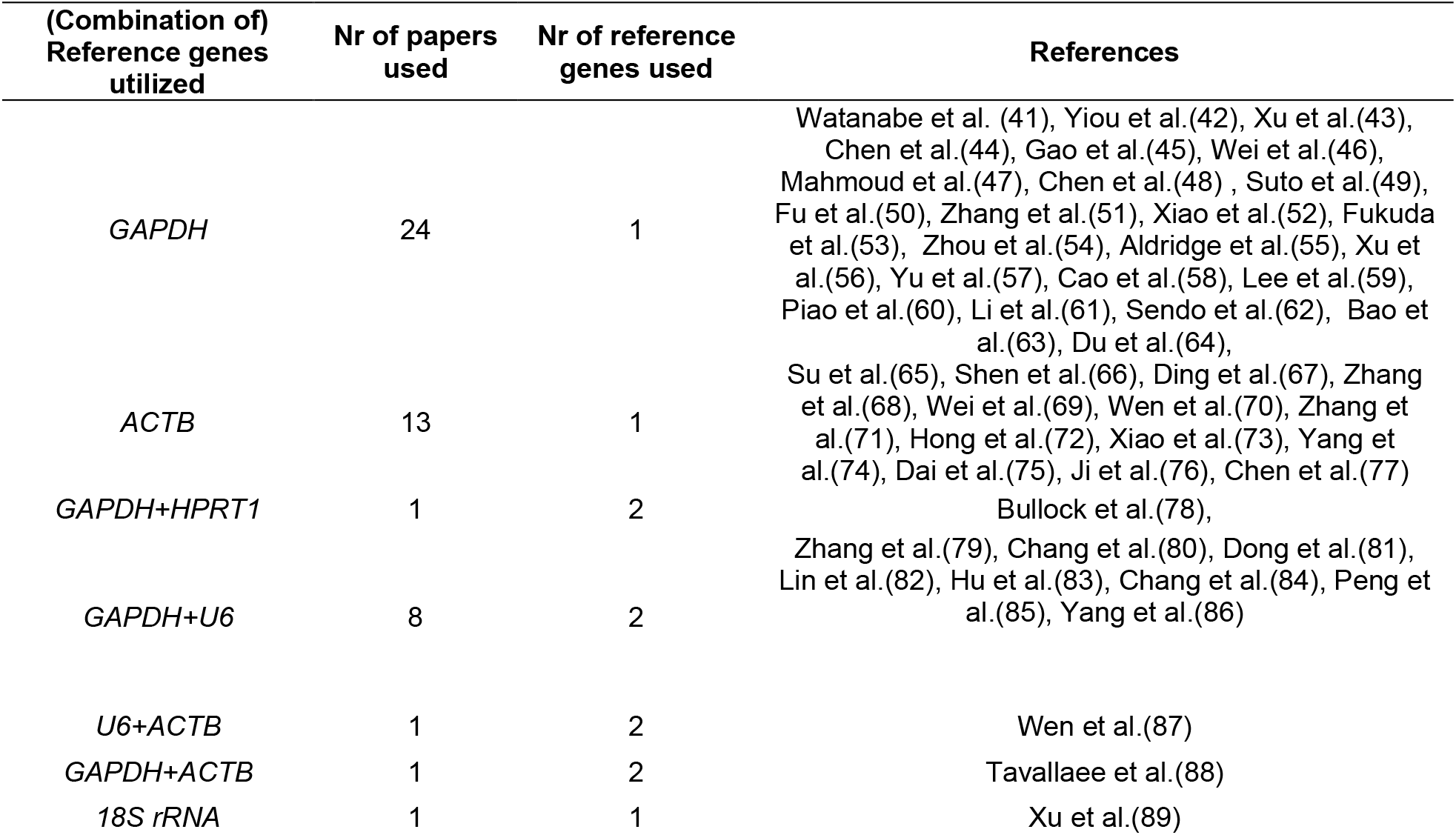
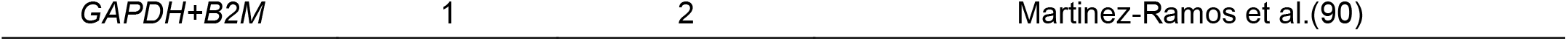
Overview of reference genes used in FLS studies published within the last 5 years.

### Step 1: Identification of candidate reference genes

First, the PubMed database was searched for research articles published within the last five years (2020-2024) that employed RT-qPCR to examine gene expression in cultured human FLS and LNSCs. Due to the lack of a sufficient number of studies on human LNSCs, this cell type was excluded from the PubMed search. To investigate the most frequently used reference genes, we randomly sampled 10 papers from each year and compiled a table summarizing the most prevalent choices (Table 2). The most frequently used reference gene for cultured FLS was *GAPDH*, followed by *ACTB*. Most related studies utilize only one reference gene as an internal control for RT-qPCR analysis. However, as previously mentioned, it is highly recommended to employ a specific number of reference genes based on the sample set and the nature of the analysis to ensure greater accuracy.

GeneVestigator is a robust tool for analysing gene expression levels in online available microarray datasets that are normalized and well annotated (5). Within GeneVestigator, we employed the ‘RefGenes’ tool to identify genes that had high expression stability across online available sample sets of FLS and LNSC microarray data. The output was checked for genes related to pathways involved in our fields of interest. Accordingly, we excluded genes related to metabolic and aging-related pathways and genes known to be related to RA pathogenesis. Moreover, the proposed candidate genes were required to have reliable expression levels within our proposed cell types.

### Step 2: Selection and expression profiles of candidate reference genes

Based on the RefGenes output, we manually selected 15 candidate reference genes. The mean Ct values and the standard deviations of the candidate reference genes are presented in Figure 3 for each transcript amplified from each biological replicate. The average Ct values ranged from 10.5 to 39.1; *18S rRNA* presented the highest and *CCD88C* transcripts presented the lowest expression levels among all the samples. Due to difficulties in designing high-quality primers for *ESX1, EIF3M* and *KCNRG*, these genes were excluded from further analysis. As a result, we identified 10 candidate reference genes for testing, in addition to two standard genes routinely used in the laboratory: *GAPDH* and *18S rRNA*.

**Figure 3.**
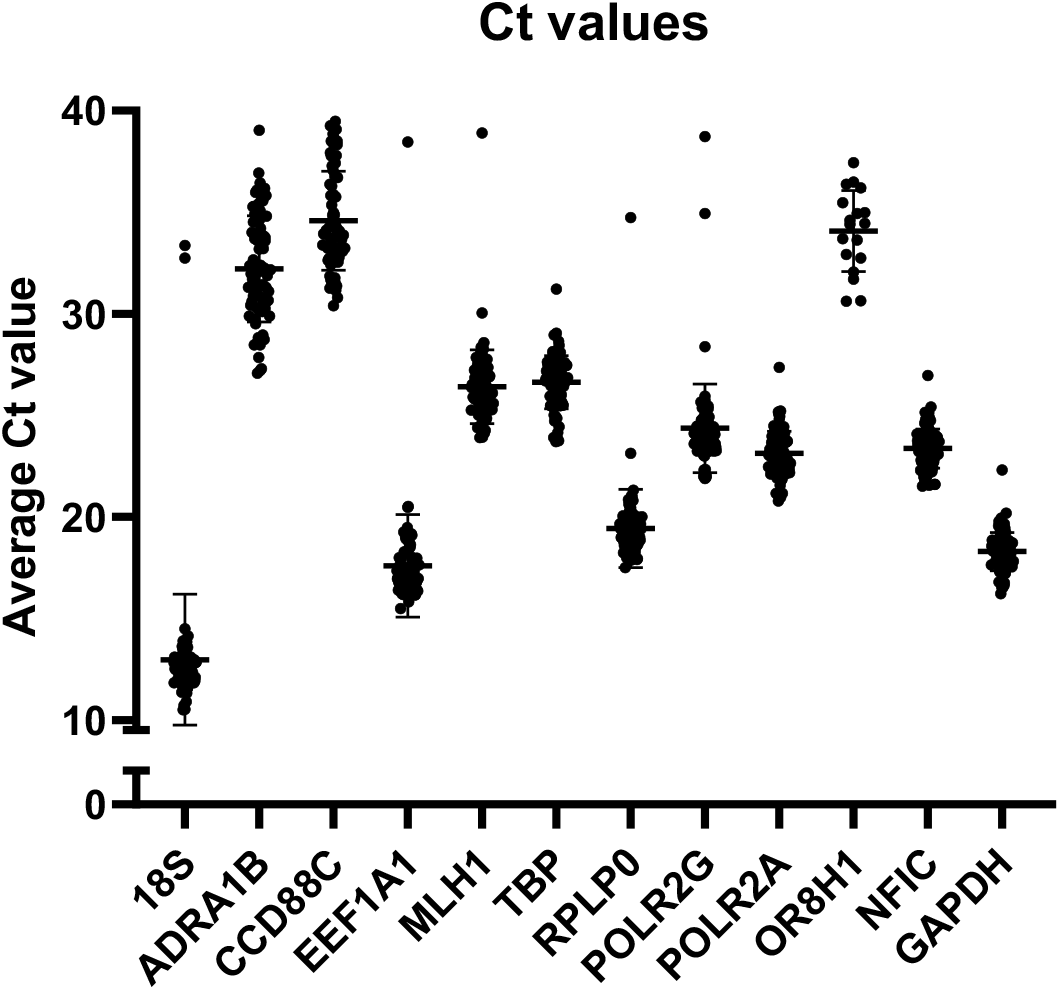
Average Ct values of untreated and treated FLS and LNSCs for each candidate reference gene. The average Ct values for all genes ranged between 10.5 and 39.1. Samples that were still undetected at qPCR cycle 40 were excluded from the analysis.

The stability of the mRNA expression levels of the selected candidate reference genes within the sample sets was assessed using the raw Ct values collected during RT-qPCR. Consequently, the RefGenes tool within GeneVestigator was used to determine the required number of reference genes in the sample sets and which candidate reference genes were stable enough to function as a reference gene within the sample sets. Next, RefFinder, which incorporates the BestKeeper, NormFinder, geNorm and delta Ct algorithms, was used to rank the candidate reference genes based on expression stability (34).

First, we performed a comprehensive analysis of all fibroblast samples in geNorm. As this did not lead to any conclusive results, the samples were grouped into different sample sets based on their tissue origin. Notably, in the LNSC sample set, the stability of the six reference genes remained consistently below the geNorm cut-off of M<0.3, irrespective of the experimental conditions. Based on the sample set, RefGenes software recommended the use of 2 reference genes as optimal for LNSCs (Figure 4). The reference genes *POLR2G* and *RPLP0* exhibited the most stable expression levels in the LNSC sample set (Figure 4, Table 3, Supplementary Table 1). Notably, *ADRA1B2* and *CCD88C* exhibited the least stable expression levels overall, with high expression variability and low expression levels.

**Figure 4.**
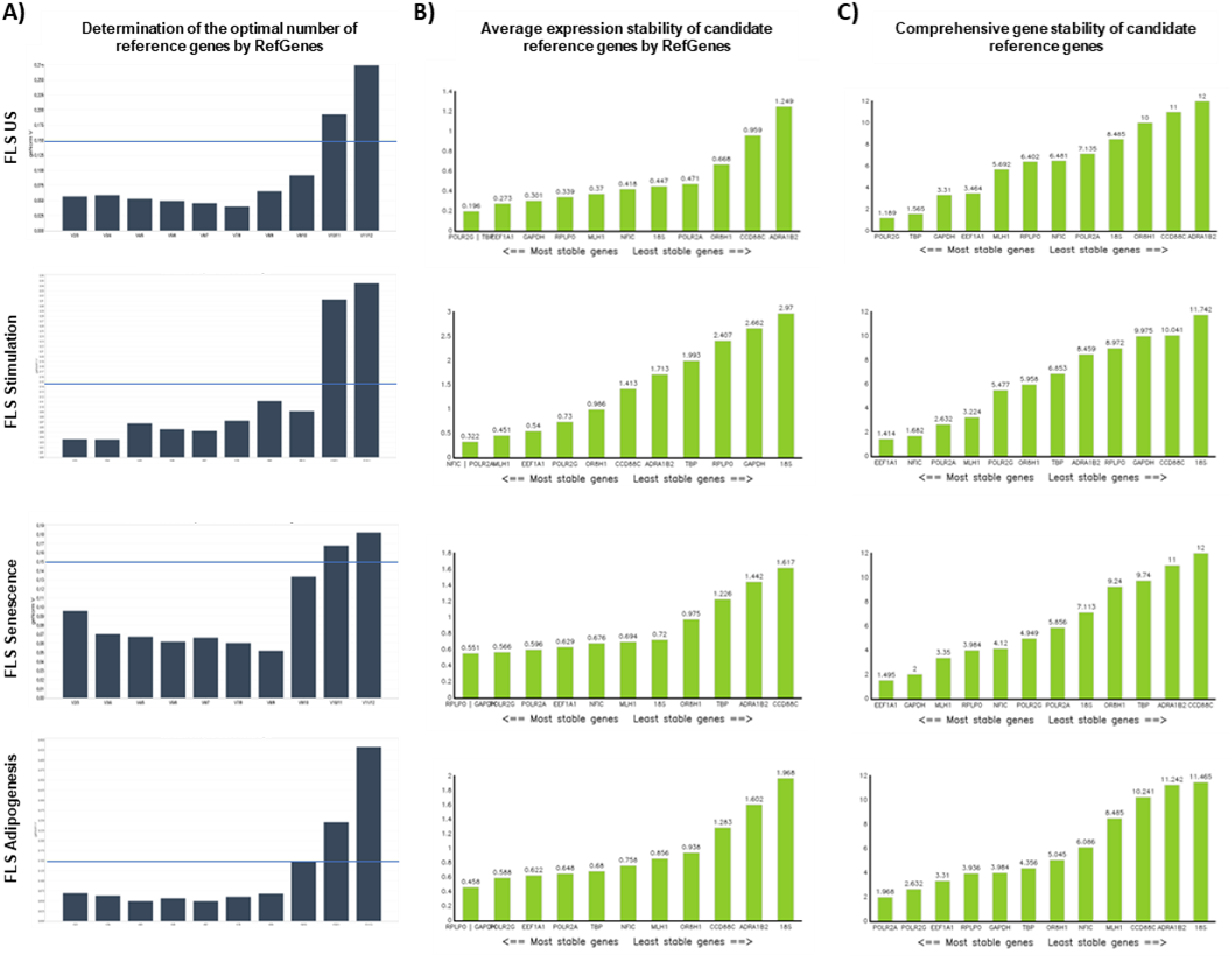
Overview of the optimal candidate reference gene number and average expression stability per candidate reference gene and experimental condition. A) Determination of the optimal number of reference genes per experimental condition. The optimal number of reference genes was determined by the V-value from the geNorm algorithm in RefGenes. B) Average expression stability of candidate reference genes defined by the M-value from the geNorm algorithm in RefGenes. C) Comprehensive gene stability of candidate reference genes defined by the overall ranking based on all algorithms implemented in RefFinder, BestKeeper, geNorm, NormFinder and the delta delta Ct method.

**Table 3.** Overview of the optimal reference genes per experimental condition and cell type.

|  | Condition | Nr of reference genes | Reference genes |
| --- | --- | --- | --- |
| FLS | US | 2 | POLR2G, TBP |
|  | Stimulation | 2 | EEF1A1, NFIC |
|  | Senescence | 2 | EEF1A1, GAPDH |
|  | Adipogenesis | 2 | POLR2A, EEF1A1* |
| LNSCs | All | 2 | POLR2G, RPLP0 |

\* The algorithms used cannot distinguish between genes from different pathways, so if multiple genes from similar pathways are included in the candidate reference gene selection, the output must be checked manually for gene function. Optimally, when more than one reference gene is recommended, every reference gene is involved in a different biological pathway. For example, the overall recommended comprehensive ranking of RefFinder recommends *POLR2A* and *POLR2G* as reference genes for the FLS adipogenesis sample set (Supplementary Table 1). These two genes are involved in the same biological pathways; therefore, it is better to use the first and third genes (*EEF1A1*) of the ranking order as a reference gene set.

However, for FLS, the results were inconsistent across the experimental conditions, with the optimal reference genes varying depending on the experimental conditions. Given the heterogeneity of reference genes for FLS, we next identified optimal reference genes for each experimental condition. Based on the RefGenes calculations for inter- and intragroup variations in the expression stability of the candidate reference genes, the algorithm pinpointed the optimal number and combination of reference genes for each of the experimental conditions used in FLS (Figure 4, Table 3, Supplementary Table 1). *POLR2G* and *TBP* were most stably expressed in the untreated FLS, whereas *EEF1A1* and *NFIC* were most stable when the cells were challenged with IFNy or TNFαLTαβ. Within the ‘senescence’ sample set, which included untreated, irradiated and dasatinib-treated FLS, *EEF1A1* and *GAPDH* were identified as the most stable reference genes. Overall, these findings underscore the importance of conducting reference gene analysis tailored to each cell type and experimental condition.

### Step 3: Benchmarking the optimal reference genes

Finally, to evaluate the recommended optimal reference gene combinations, the relative expression levels of metabolism-related genes in FLS were evaluated. Gene expression normalization was performed using the selected optimal reference genes *POLR2G* and *TBP* and the traditionally used reference gene *18S rRNA*. Therefore, the values of each target gene were either normalized to the geometric mean expression levels of the two reference genes (Figure 5) or to the expression levels of *18S rRNA* alone (Figure 5). The observed relative expression levels showed different expression patterns between the control, RA-risk, RA, and OA samples depending on the reference gene(s) used. This notable difference between the two normalization methods underscores the critical importance of accurate reference gene selection in ensuring reliable gene expression analysis.

**Figure 5.**
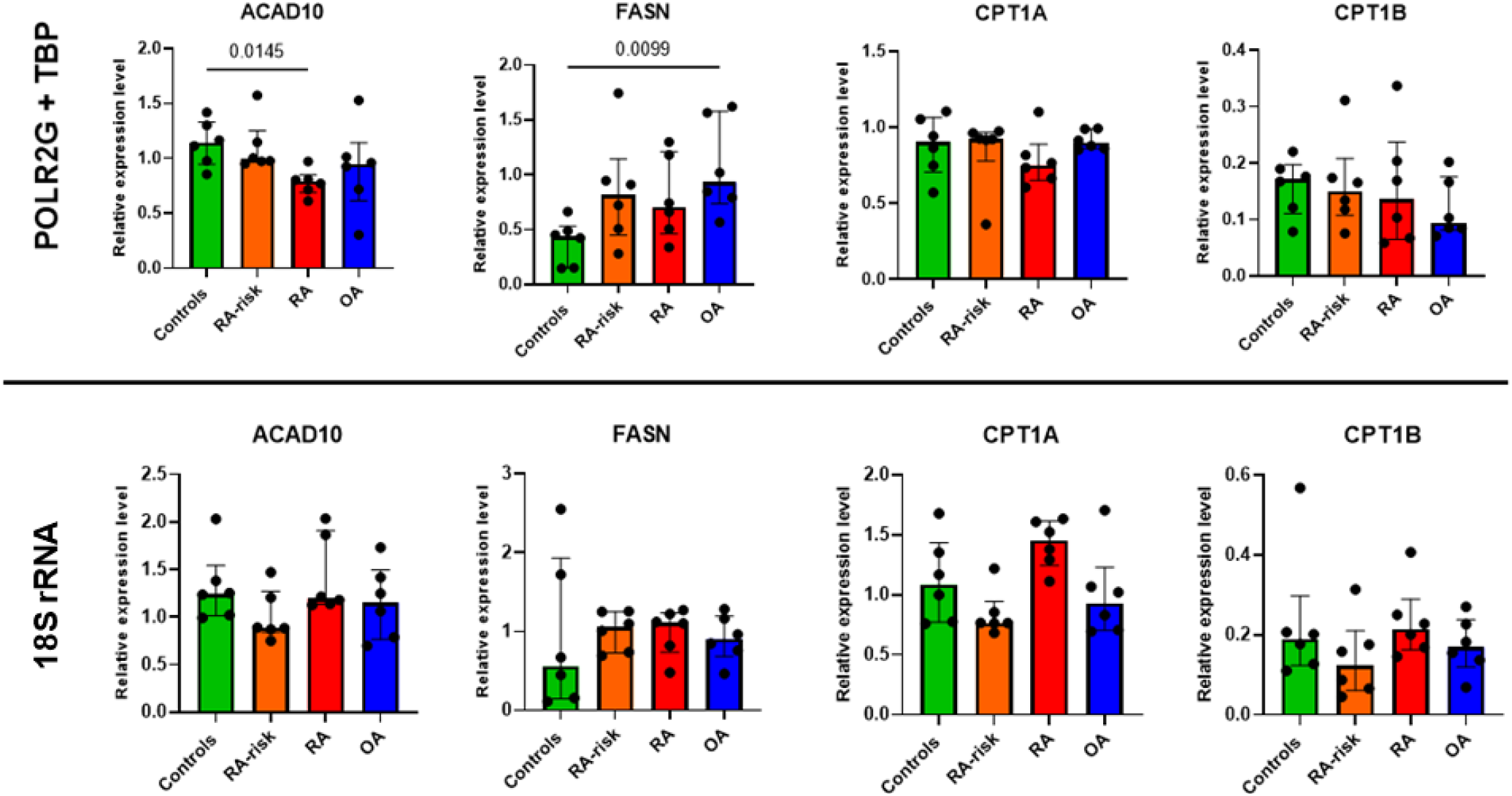
Relative expression levels of metabolic genes in FLS normalized by optimal and nonoptimal reference genes. Relative mRNA expression levels of ACAD10, FASN, CPT1A and CPT1B. All donors passage 4, n=6 per group. Data are presented as median + interquartile range, and significant differences were determined using Kruskal-Wallis + Dunn’s multiple comparisons test.

## Discussion

The selection of suitable reference genes is crucial for ensuring the accuracy of RT–qPCR data normalization. A suitable reference gene should exhibit stable mRNA expression across all cell types and experimental conditions of interest for comparison (5). No single universally stable and reliable reference gene exists. Therefore, the identification of optimal reference genes should be performed for each specific comparison of interest due to variability in gene expression levels between different tissues, cell types and exposures (3).

Caution is warranted because no gene exhibits universal stability, and commonly used reference genes often yield high transcript abundances compared to target genes of interest (2). For each specific biological sample, different genes might remain consistently stable within a specific experimental context. These genes can differ from commonly used reference genes, which are selected based on their presumed stability across a wide range of conditions and samples (5). Therefore, the stability of (reference) genes is unique to the specific biological context being studied rather than being universally stable in all experimental settings. This highlights the importance of identifying and using sample-specific stable reference genes for accurate gene expression analysis. Therefore, to enhance the precision of calculations and normalization, it is advisable to employ multiple reference genes rather than relying on a single reference gene (8).

Our research focused on investigating the stromal microenvironment of RA patients, with a specific emphasis on FLS and LNSCs. Previous studies have not validated reference genes for transcript normalization in these cell populations from RA patients. Given the crucial role of fibroblasts in RA, identifying reliable reference genes for accurate gene expression analysis is essential. In this study, we evaluated the expression stability of candidate reference genes in fibroblasts from RA and OA patients, seronegative controls and individuals at risk of developing RA under different experimental conditions (Figure 6). By employing our optimized strategy, we determined that for both FLS and LNSCs, it is best to analyse every experimental sample using the geometric mean of two reference genes to ensure accurate normalization. Interestingly, for LNSCs, *POLR2G* and *RPLP0* were the most stable across all the experimental conditions tested, while for FLS, the most stable reference genes varied for each experimental condition. Validation of the recommended reference gene set in a cohort of untreated FLS showed that the expression profiles of several target genes highly varied depending on the reference gene(s) used. This underscores the value of this approach in enhancing the reliability of gene expression analysis.

**Figure 6.**
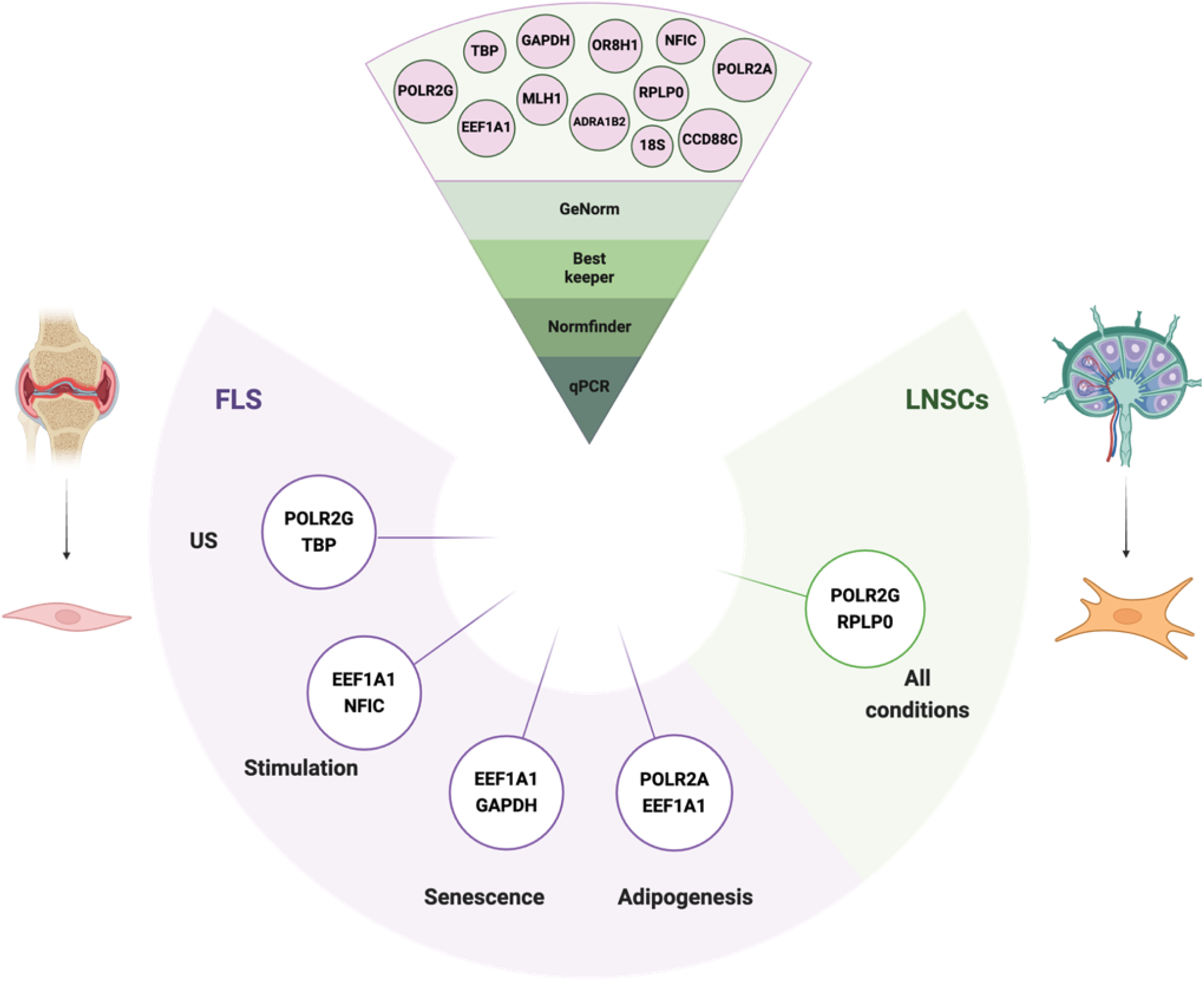
Schematic representation of the overall experimental design and results. The top part shows the 12 candidate reference genes selected based on the RefGenes results. The qPCR data of these genes, which were acquired from different cell types and under different experimental conditions, were evaluated using software tools such as geNorm, BestKeeper, and NormFinder to identify the most stable genes for each experimental condition. Finally, the stability of these genes was validated in a complete donor cohort using qPCR. We identified two reference genes per experimental condition, indicated in green for LNSCs and purple for FLS.

When defining optimal reference genes for an experiment, a few points must be considered. These include the presence of pseudogenes, biased reference gene recommendations due to limited software datasets in the available software, and the time-consuming nature of performing and analysing the data. Pseudogenes, nonfunctional copies of genes caused by mutations, have sequences similar to those of functional genes (41). Consequently, when using reference genes, the sequences from pseudogenes can result in nonspecific amplification during PCR. This can lead to inaccuracies in measuring gene expression and make it challenging to interpret experimental results accurately. Therefore, genes with many pseudogenes should be avoided as much as possible. Furthermore, the variability in experimental conditions across different studies necessitates the careful selection of reliable reference genes. Datasets in software tools such as GeneVestigator may not encompass all experimental treatments, potentially leading to biased selection of reference gene candidates. Nonetheless, it is worth noting that the available datasets used in this manuscript are extensive. Despite these limitations, this approach offers significant advantages. This approach provides a more robust alternative to relying on single gene markers for qPCR analyses, especially considering the variability across different treatments and cell types.

In conclusion, we recommend employing *POLR2G* and *RPLP0* as reference genes when examining gene expression variations among LNSCs across all exposures investigated in this study. Given the diversity of responses observed in FLS, we suggest employing different reference genes depending on the specific treatment conditions. Specifically, for untreated FLS, we advocate for the use of *POLR2G* and *TBP*, while for FLS subjected to IFNγ or TNFαLTαβ stimulation, we propose the use of *EEF1A1* and *NFIC*. Furthermore, for investigating senescence in FLS, *EEF1A1* and *GAPDH* are the most reliable reference genes. Finally, for studies concerning FLS undergoing adipogenesis, we suggest the use of *POLR2A* and *EEF1A1*. It is important to note, however, that while these recommendations are based on our specific set of cell lines and treatments, extrapolating these findings to other cell lines and conditions requires rigorous experimental validation. These findings highlight the necessity for meticulous consideration of reference genes in experimental design, ensuring robustness and accuracy in gene expression studies. Overall, this study provides a strategy to select and validate potential candidate reference genes to normalize RT–qPCR results according to sample type and experimental conditions of interest, ensuring the accuracy of gene expression measurements.

