## Supplemental Table 1 for "Identification of stable reference genes for quantitative real-time PCR in human fibroblasts from lymph nodes and synovium"

### Supplementary data

**Supplementary Table 1. Overview of the ranking order of the candidate reference genes per experimental condition**

| | | Ranking Order (Better–Good–Average) | | | | | | | | | | | |
|---|---|---|---|---|---|---|---|---|---|---|---|---|---|
| US | Method | 1 | 2 | 3 | 4 | 5 | 6 | 7 | 8 | 9 | 10 | 11 | 12 |
| | Delta Ct | POLR2G | TBP | GAPDH | EEF1A1 | MLH1 | RPLP0 | NFIC | 18S | POLR2A | OR8H1 | CCD88C | ADRA1B2 |
| | BestKeeper | TBP | POLR2G | EEF1A1 | POLR2A | GAPDH | NFIC | MLH1 | RPLP0 | 18S | OR8H1 | CCD88C | ADRA1B2 |
| | NormFinder | POLR2G | GAPDH | TBP | EEF1A1 | MLH1 | NFIC | RPLP0 | POLR2A | 18S | OR8H1 | CCD88C | ADRA1B2 |
| | GeNorm | POLR2G \| TBP | | EEF1A1 | GAPDH | RPLP0 | MLH1 | NFIC | 18S | POLR2A | OR8H1 | CCD88C | ADRA1B2 |
| | Rec comprehen ranking | POLR2G | TBP | GAPDH | EEF1A1 | MLH1 | RPLP0 | NFIC | POLR2A | 18S | OR8H1 | CCD88C | ADRA1B2 |

| | | Ranking Order (Better–Good–Average) | | | | | | | | | | | |
|---|---|---|---|---|---|---|---|---|---|---|---|---|---|
| Senescence | Method | 1 | 2 | 3 | 4 | 5 | 6 | 7 | 8 | 9 | 10 | 11 | 12 |
| | Delta Ct | EEF1A1 | GAPDH | MLH1 | NFIC | POLR2G | RPLP0 | POLR2A | 18S | OR8H1 | TBP | ADRA1B2 | CCD88C |
| | BestKeeper | EEF1A1 | MLH1 | NFIC | GAPDH | 18S | RPLP0 | POLR2A | POLR2G | TBP | OR8H1 | ADRA1B2 | CCD88C |
| | NormFinder | EEF1A1 | GAPDH | MLH1 | NFIC | POLR2G | POLR2A | RPLP0 | 18S | OR8H1 | TBP | ADRA1B2 | CCD88C |
| | GeNorm | RPLP0 \| GAPDH | | POLR2G | POLR2A | EEF1A1 | NFIC | MLH1 | 18S | OR8H1 | TBP | ADRA1B2 | CCD88C |
| | Rec comprehen ranking | EEF1A1 | GAPDH | MLH1 | RPLP0 | NFIC | POLR2G | POLR2A | 18S | OR8H1 | TBP | ADRA1B2 | CCD88C |

| | | Ranking Order (Better–Good–Average) | | | | | | | | | | | |
|---|---|---|---|---|---|---|---|---|---|---|---|---|---|
| Adipogenesis | Method | 1 | 2 | 3 | 4 | 5 | 6 | 7 | 8 | 9 | 10 | 11 | 12 |
| | Delta Ct | POLR2A | POLR2G | EEF1A1 | TBP | RPLP0 | GAPDH | NFIC | MLH1 | OR8H1 | CCD88C | ADRA1B2 | 18S |
| | BestKeeper | OR8H1 | EEF1A1 | POLR2A | POLR2G | TBP | GAPDH | NFIC | RPLP0 | MLH1 | 18S | CCD88C | ADRA1B2 |
| | NormFinder | POLR2A | POLR2G | TBP | NFIC | EEF1A1 | RPLP0 | GAPDH | OR8H1 | MLH1 | CCD88C | ADRA1B2 | 18S |
| | GeNorm | RPLP0 \| GAPDH | | POLR2G | EEF1A1 | POLR2A | TBP | NFIC | MLH1 | OR8H1 | CCD88C | ADRA1B2 | 18S |
| | Rec comprehen ranking | POLR2A | POLR2G | EEF1A1 | RPLP0 | GAPDH | TBP | OR8H1 | NFIC | MLH1 | CCD88C | ADRA1B2 | 18S |

| | | Ranking Order (Better–Good–Average) | | | | | | | | | | | |
|---|---|---|---|---|---|---|---|---|---|---|---|---|---|
| Stimulation | Method | 1 | 2 | 3 | 4 | 5 | 6 | 7 | 8 | 9 | 10 | 11 | 12 |
| | Delta Ct | EEF1A1 | NFIC | MLH1 | POLR2A | POLR2G | OR8H1 | TBP | ADRA1B2 | RPLP0 | GAPDH | CCD88C | 18S |
| | BestKeeper | EEF1A1 | NFIC | POLR2A | MLH1 | OR8H1 | POLR2G | TBP | RPLP0 | GAPDH | ADRA1B2 | 18S | CCD88C |
| | NormFinder | EEF1A1 | NFIC | MLH1 | POLR2A | TBP | POLR2G | OR8H1 | ADRA1B2 | RPLP0 | GAPDH | CCD88C | 18S |
| | GeNorm | NFIC \| POLR2A | | MLH1 | EEF1A1 | POLR2G | OR8H1 | CCD88C | ADRA1B2 | TBP | RPLP0 | GAPDH | 18S |
| | Rec comprehen ranking | EEF1A1 | NFIC | POLR2A | MLH1 | POLR2G | OR8H1 | TBP | ADRA1B2 | RPLP0 | GAPDH | CCD88C | 18S |

| | | Ranking Order (Better–Good–Average) | | | | | | | | | | | |
|---|---|---|---|---|---|---|---|---|---|---|---|---|---|
| All LNSC conditions | Method | 1 | 2 | 3 | 4 | 5 | 6 | 7 | 8 | 9 | 10 | 11 | 12 |
| | Delta CT | RPLP0 | POLR2G | EEF1A1 | POLR2A | GAPDH | NFIC | 18S | OR8H1 | MLH1 | TBP | ADRA1B2 | CCD88C |
| | BestKeeper | RPLP0 | POLR2G | EEF1A1 | OR8H1 | MLH1 | POLR2A | GAPDH | NFIC | TBP | 18S | ADRA1B2 | CCD88C |
| | Normfinder | EEF1A1 | RPLP0 | 18S | POLR2G | POLR2A | GAPDH | NFIC | OR8H1 | MLH1 | TBP | ADRA1B2 | CCD88C |
| | Genorm | GAPDH \| POLR2A | | POLR2G | NFIC | RPLP0 | EEF1A1 | 18S | OR8H1 | MLH1 | TBP | ADRA1B2 | CCD88C |
| | Rec comprehen ranking | RPLP0 | POLR2G | EEF1A1 | POLR2A | GAPDH | NFIC | 18S | OR8H1 | MLH1 | TBP | ADRA1B2 | CCD88C |

Rec comprehen ranking = recommended comprehensive ranking.
